# Characterization of the gut DNA and RNA viromes in a cohort of Chinese residents and visiting Pakistanis

**DOI:** 10.1101/2020.07.28.226019

**Authors:** Qiulong Yan, Yu Wang, Xiuli Chen, Hao Jin, Guangyang Wang, Kuiqing Guan, Yue Zhang, Pan Zhang, Taj Ayaz, Yanshan Liang, Junyi Wang, Guangyi Cui, Yuanyuan Sun, Manchun Xiao, Aiqin Zhang, Peng Li, Xueyang Liu, Yufang Ma, Shenghui Li, Tonghui Ma

## Abstract

**Background:** Trillions of viruses inhabit the gastrointestinal tract. Some of them have been well-studied on their roles in infection and human health, but the majority remain unsurveyed. It has been established that the composition of the gut virome is highly variable based on the changes of diet, physical state, and environmental factors. However, the effect of host genetic factors, e.g. ethnic origin, on the gut virome is rarely investigated.

**Methods and Results:** Here, we characterized and compared the gut virome in a cohort of local Chinese residents and visiting Pakistani individuals, each group containing 24 healthy adults and 6 children. Using metagenomic shotgun sequencing and assembly of fecal samples, a huge number of viral operational taxonomic units (vOTUs) were identified for profiling the DNA and RNA viromes. National background contributed a primary variation to individuals’ gut virome. Compared with the Chinese adults, the Pakistan adults showed higher macrodiversity and different compositional and functional structures in their DNA virome and lower diversity and altered composition in their RNA virome. The virome variations of Pakistan children were inherited from the that of the adults but also tended to share similar characteristics with the Chinese cohort. We also analyzed and compared the bacterial microbiome between two cohorts and further revealed numerous connections between virus and bacterial host. Statistically, the gut DNA and RNA viromes were covariant to some extent (*p*<0.001), and they both influenced the holistic bacterial composition and vice versa.

**Conclusions:** This study provides an overview of gut viral community in Chinese and visiting Pakistanis and proposes a considerable role of ethnic origin in shaping the virome.

## Background

The human gut is a large reservoir of microorganisms, containing 10^11^-10^12^ bacterial cells [1, 2], 10^9^-10^12^ viral particles [3, 4], and small quantities of archaea and eukaryotes in per gram of feces [5]. Benefiting from the development of high throughput sequencing techniques (e.g. amplicon or whole-metagenomic sequencing), the gut bacterial community have been well studied over the past years [6-8]. Gut bacteria was shown to exert profound effects on regulating host metabolism [9, 10], and thereby had been linked to host health and diseases [11, 12]. However, as another part of the gut microbial ecosystem, the holistic viral community of enteric microbiome (or “gut virome”) was less well characterized [13]. Virus has a very flexible small genome ranging from a few to several hundred kilobases [14], which corresponds to approximately 1% of the bacterial genome (in average, 2-4 Mbp) [15, 16]. The gut virome was predominantly composed of two taxa of bacteriophages, double-stranded DNA *Caudovirales* and single-stranded DNA *Microviridae*, which constituted over 80% relative abundance of viral populations in human intestine [17]. The *crAssphage* and *crAss-like* phages, a type of *Caudovirales* members that characteristically infect *Bacteroides* spp., represented the highest abundance in healthy human gut [18, 19]. In addition to bacteriophages, eukaryotic viruses, archaeal viruses, and RNA viruses were also important components of gut virome [20, 21].

Due to the limitation of viral abundance in human gut, routine whole-metagenomic sequencing of fecal microbiome can produce only a small proportion of viral sequences for further analysis. Recently, virus-like particle (VLP) enrichment and subsequently metagenomic sequencing provided a prospective application for fully delineating the gut virome [22, 23]. Based on the VLP technique, studies had showed that the normal gut virome was partly inherited from mother [24, 25], potentially transferred between twins [4], and continuously expanded during the first years of life [21]. In addition, longitudinal analysis revealed that the gut virome of healthy adults was highly diverse, temporally stable, and individually specific [14]. Disease-induced alterations of the gut virome had also be reported in multiple gastrointestinal and systemic disorders, including colorectal cancer [26, 27], inflammatory bowel disease [17, 28], type I diabetes [29], and coronary heart disease [30]. These studies suggest a significant role of gut virome in human health, however, some essential issues of human gut virome, such as population heterogeneity and impacts of geography, lifestyle or environment, is still in shortage.

By studying the gut microbiome of migrated or short-term visiting peoples, previous studies had shown that their microbiota was markedly remodeled upon environmental change, but yet accompanied with maintenance of numerous individual or ethnic microbial characteristics [31-34]. Herein, we depicted the compositional differences of gut virome between Chinese residents (n = 24) and visiting Pakistani (n = 24) individuals living in the same city and also examined the repeatability of these differences in their child offsprings (respective n = 6). We quantified the DNA and RNA viromes from fecal VLPs, and parallelly measured the bacterial microbiome for virus-bacteria association analysis. This pilot study provided evidences for the effect of ethnic backgrounds on human gut virome.

## Results

### Population characteristics and study design

This study included 30 Chinese residents and 30 visiting Pakistani individuals who were recruited at Dalian Medical University in March 2019. Both cohorts consisted of 24 healthy adults and 6 of their child offsprings (**Table 1**). All adults were students or young teachers of the Dalian Medical University, and the Pakistani adults and children had arrived in China for 0-18 months (average of 11 months) and 0-15 months (average of 9 months), respectively. Notably, the Chinese and Pakistani adults showed significant differences on their body mass index (BMI), dietary habit, and drinking and smoking rates (**Table 1; Table S1**), which seemed to be due to ethnic and lifestyle differences.

**Table 1.** Characteristics of the subjects.

|  | Adults |  |  | Children |  |  |
| --- | --- | --- | --- | --- | --- | --- |
|  | Chinese | Pakistani | *P*-value | Chinese | Pakistani | *P*-value |
| Number of subjects | 24 | 24 |  | 6 | 6 |  |
| Sex, F/M | 1/23 | 1/23 | 1.000 | 3/3 | 3/3 | 1.000 |
| Age, years | 26.0±4.3 | 29.1±3.7 | 0.011 | 2.8±1.8 | 2.8±1.7 | 1.000 |
| Weight, kg | 69.6±11.0 | 76.7±15.6 | 0.076 | 14±5.4 | 13.3±4.2 | 0.794 |
| BMI, kg/m<sup>2</sup> | 22.8±2.8 | 25.6±4.5 | 0.011 | 16.0±2.0 | 17.4±3.1 | 0.396 |
| Drinking, % | 50% | 8.3% | 0.003 | 0% | 0% | 1.000 |
| Smoking, % | 16.7% | 33.3% | 0.030 | 0% | 0% | 1.000 |
| Antibiotics (≤2mons), % | 8.3% | 8.3% | 1.000 | 0% | 0% | 1.000 |
| Prebiotics (≤2mons), % | 58.3% | 41.7% | 0.387 | 66.7% | 50% | 1.000 |
| Living in China, mons |  | 11±4 |  |  | 9±6 |  |

The data for age, weight, and BMI were presented as mean ± sd. *P*-values for age, weight, and BMI were calculated by Student's t-test, and for sex, drinking, smoking, antibiotics, and prebiotics were calculated by Fisher's exact test.

Fecal samples of all participants were collected and treated using a unified approach (see Methods). To depict the gut viral characteristics of healthy individuals, we extracted DNA and RNA from fecal VLP factions and performed high throughput shotgun sequencing using the Illumina platform. To extend the content of total microbial community, the bacterial microbiome of feces was also profiled using whole-metagenomic sequencing. The analytical workflow of the DNA virome, RNA virome, and bacterial microbiome was shown in **Figure 1**. Focusing on the comparison of gut viromes between Chinese and Pakistani individuals, overall, this study included six sections to elaborate the results:

1-2. DNA virome and its functional characteristics.

3-4. RNA virome and the concordance between DNA and RNA viromes.

5-6. Bacterial microbiome and the virus-bacteria associations.

**Figure 1.**
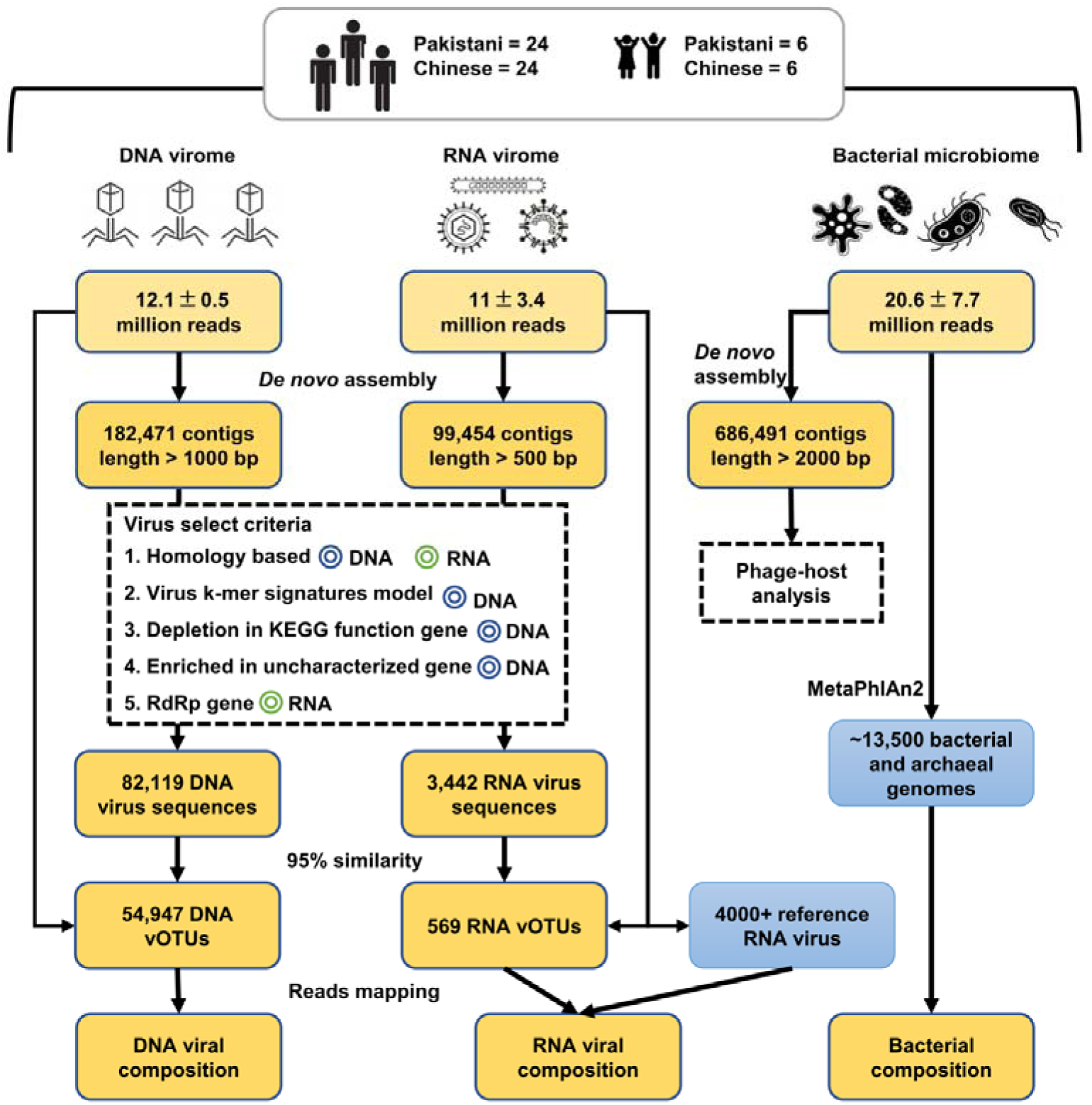
Overview of the workflow for analyzing of DNA virome, RNA virome, and bacterial microbiome.

### Comparison of DNA viral community

We obtained 782 million high-quality non-human reads (12.1 ± 0.5 million per sample) through shotgun sequencing of the DNA viral community of 60 fecal samples. The reads were *de novo* assembled into 182,471 contigs with the minimum length threshold of 1kbp, of which 45.0% (82,119) were recognized as highly credible viral fragments based on their sequence features and homology to known viral genomes (**Figure 1**). The remaining contigs were from bacterial or eukaryotic contaminations (26.8%) and dependency-associated sequences (6.0%), and 22.2% contigs were still unclassifiable. Despite that, average 82.3% of sequencing reads in all samples were captured by the viral contigs, revealing well representativeness of the high-abundance viral contents in human gut DNA virome. The viral contigs were further clustered into 54,947 “viral operational taxonomic units (vOTUs)” (a phylogenetic definition of discrete viral lineage that corresponds to “species” in prokaryotes, also named “viral population” [35]) by removing the redundant contigs of 95% nucleotide similarity. These vOTUs represented an average size of 3,054 ± 2,868 bp (**Figure S1**), which was comparable with similar studies [14] but remarkable lower than that of the available viral genomes (average 38.5 kbp for ∼6,500 complete virus isolates from the RefSeq database), suggesting that the vOTUs were mostly fragmented genomes. Only 33.6% of vOTUs could be annotated into specific family, highlighting a considerable novelty of gut virome.

Rarefaction analysis showed that, despite the rarefaction curve was unsaturated under current number of samples in each group, the vOTU richness was significantly higher in Pakistani adults than in Chinese adults (*p*=0.008, **Figure 2a**). The within-sample diversity pattern of gut DNA viromes was assessed by macrodiversity (Shannon index) and microdiversity (nucleotide diversity or π [35]) at the vOTU level. The Chinese adults showed a lower Shannon index than the Pakistani adults, similarly for the children (**Figure 2b**), but no significant difference in microdiversity was detected between Chinese and Pakistanis (**Figure 2c**).

**Figure 2.**
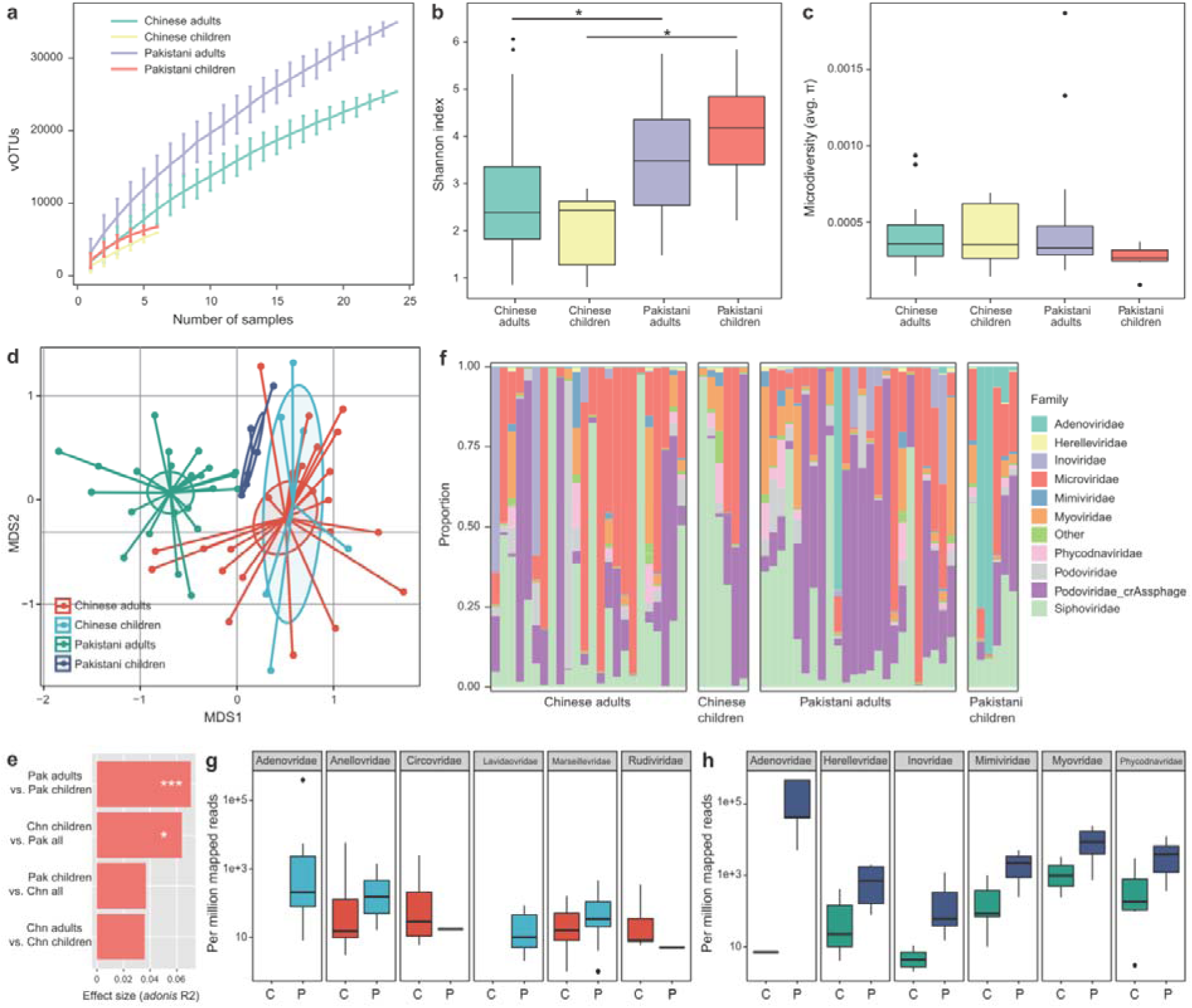
Differences in gut DNA virome between Chinese and Pakistanis. **a**, Rarefaction curve analysis of number of vOTUs on each group of samples. The number of identified vOTUs in different groups is calculated based on a randomly selected specific number of samples with 30 replacements, and the median and quartiles numbers are plotted. **b-c**, Boxplot shows the macrodiversity **(b)** and microdiversity **(c)** that differ among four groups. The significance level in the Student’s t test is denoted as: *, *q*<0.05; **, *q*<0.01. **d**, NMDS analysis based on the composition of virome, revealing the separations between different groups. The location of samples (represented by nodes) in the first two multidimensional scales are shown. Lines connect samples in the same group, and circles cover samples near the center of gravity for each group. **e**, PERMANOVA analysis reveals that the virome of Pakistani children are similar with the Chinese subjects (*adonis p*>0.05). The effect sizes and *p*-values of the *adonis* analysis are shown. **f**, Composition of gut virome at the family level. **g-h**, Boxplot shows the differential viral families of adults **(g)** and children **(h)** when compared between Chinese and Pakistanis. C, Chinese individuals; P, Pakistani individuals. For boxplot, boxes represent the interquartile range between the first and third quartiles and median (internal line); whiskers denote the lowest and highest values within 1.5 times the range of the first and third quartiles, respectively; and nodes represent outliers beyond the whiskers.

Next, we undertook a non-metric multidimensional scaling (NMDS) analysis to further understand the differences in fecal DNA viral communities between Chinese and Pakistanis. Clear separations were revealed in the viromes of both adults and children between Chinese and Pakistanis (*adonis p*<0.001 for both adults and children; **Figure 2d**). Notably, we also found that 1) the viral communities of Chinese adults and children were similar, but those of Pakistani adults and children were differed, and 2) the viral communities of Pakistani children were closer to Chinese subjects when compared with those of Pakistani adults. These findings were validated by the permutational multivariate analysis of variance (PERMANOVA) (**Figure 2e**).

We finally compared the DNA virome composition of Chinese and Pakistani at the family level, ignoring the family-level unclassified vOTUs (which represented only 33.1% of total sequences). The most dominant viral families in all samples were *Podoviridae*-crAssphage (average relative abundance, 27.0 ± 30.7%), *Siphoviridae* (24.8 ± 25.5%) and *Adenoviridae* (23.7 ± 28.1%) (**Figure 2f**). Compared with the Chinese adults, the viral communities of the Pakistani adults showed a significant increase of *Adenoviridae, Anelloviridae, Marseilleviridae*, and *Lavidaviridae*, and a remarkable depletion of *Circoviridae* and *Rudiviridae* (Mann-Whitney U test, *q*<0.05; **Figure 2g**). *Adenoviridae, Myoviridae, Phycodnaviridae, Mimiviridae, Herelleviridae*, and *Inoviridae* were significant higher in viral communities of Pakistani children (**Figure 2h**), as compared with the Chinese children, while no viral family was lower.

### Functional analysis of DNA virome

To better elucidate the functional capacity of the DNA viromes, we predicted a total of 221,418 protein-coding genes from the vOTUs (average of 4 genes per vOTU) and annotated functions of 24.2% of these genes based on the KEGG (Kyoto Encyclopedia of Genes and Genomes) [36] database. Analysis on KEGG pathway level B showed that functions involved in genetic information procession and signal and cellar processes are dominant in all samples (**Figure 3a**), suggesting that these are core functions of the gut DNA virome. Compared with the Chinese adults, viral functions in the Pakistani adults were significantly decreased involving “protein families: metabolism”, amino acid metabolism, antimicrobial drug resistance, cell motility, and substance dependence, and increased in immune disease (Mann-Whitney U test, *q*<0.05; **Figure 3b**). For example, a putative hemolysin enzyme (K03699) that encoded by several *Myoviridae* and *Siphoviridae* viruses showed over 10-fold enrichment in the virome of Chinese adults compared to that of Pakistani adults. When compared with the Chinese children, a number of important functions, including carbohydrate metabolism, signal transduction, and cell growth and death, were significantly higher in the viral communities of Pakistani children, while the “protein families: genetic information processing” were lower (**Figure 3c**).

**Figure 3.**
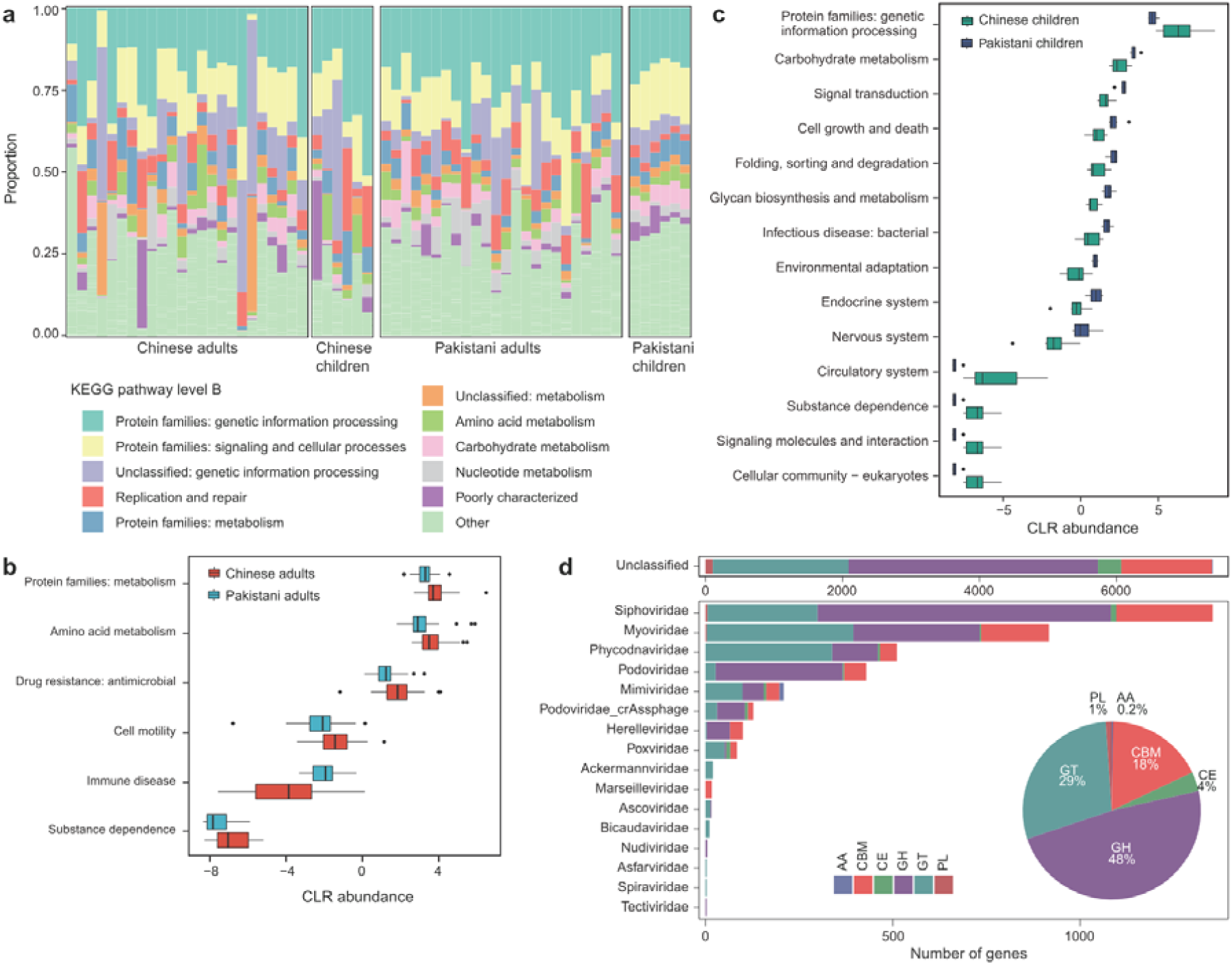
Comparison of DNA viral functions between Chinese and Pakistanis. **a**, Composition of viral functional categories at the KEGG pathway level B. **b-c**, Boxplot shows the KEGG pathways that differed in abundance between Chinese adults and Pakistani adults **(b)** and between Chinese children and Pakistani children **(c)**. Boxes represent the interquartile range between the first and third quartiles and median (internal line); whiskers denote the lowest and highest values within 1.5 times the range of the first and third quartiles, respectively; and nodes represent outliers beyond the whiskers. **d**, The taxonomic distribution of CAZymes. GH, glycoside hydrolase; GT glycosyl transferase; CBM, carbohydrate binding; CE, carbohydrate esterase; PL, polysaccharide lyase; AA auxiliary activity.

We identified a total of 11,242 CAZymes (Carbohydrate-active enzymes [37]) from the viral genes, including 5,437 glycoside hydrolases, 3,270 glycosyl transferases, 1,993 carbohydrate binding, 396 carbohydrate esterases, 120 polysaccharide lyases, and 26 auxiliary activities (**Figure 3d**). The majority (65.9%) of CAZymes were encoded by unclassified vOTUs, followed by *Siphoviridae* (12.1%) and *Myoviridae* (8.2%), suggesting their important roles in carbohydrate metabolism in gut viral ecosystem. Moreover, we also identified 37 acquired antibiotic resistance genes (ARGs) from the DNA vOTUs (**Table S2**). Most of these ARGs were related to tetracycline resistance (n = 12), macrolide resistance (n = 7), beta-lactamase (n = 7), and aminoglycoside resistance (n = 6). Taken together, these findings revealed that the DNA virus can widely express the carbohydrate metabolism-associated genes and are potentially involved into carrying and transmission of antibiotic resistance genes.

### Comparison of RNA viral community

For RNA virome, we performed shotgun metatranscriptomic sequencing of 60 fecal samples described above and obtained 671 million reads (11 ± 3.4 million per sample) after removing the low-quality reads and bacterial ribosomal RNA contamination. A total of 99,454 contigs with minimum length threshold of 500 bp were assembled, 3,442 (3.5%) of which were identified as highly credible RNA viral fragments via blasting against the available RNA viral genomes and searching of the RNA-dependent RNA polymerase (RdRp) sequences (**Figure 1**). 25.4% of these RNA viruses contained at least one RdRp gene, while 28 viral RdRp genes had no homology with any known virus in NCBI database. We obtained 569 RNA vOTUs based on clustering at 95% nucleic acid level similarity. The average size of these vOTUs was 1,162 ± 916 bp, which was fragmented compared with the available RNA viral genomes (average 7.4 kbp from ∼4,000 isolates). Furthermore, considering that only average 24.8% reads of all samples were covered from the RNA vOTUs, we also used the available RNA viral genomes from the RefSeq database as a reference for analyzing of the gut RNA virome. 118 available RNA viruses were observed in our samples, which covered additional 1.3% reads (in average) for further analysis. Rarefaction analysis showed that the detection of RNA virus was increased with the number of samples, and the accumulative curve was nearly saturated at nearly 10 samples (**Figure 4a**). This is due to our RNA virus pipeline mainly focused on the known species and the sequence containing a RdRp gene, but high proportions of virus remain untagged and many of them are independent on RdRp gene [38]. Compared with Pakistanis, the macrodiversity (Shannon index) was significantly higher in Chinese adults, but there was no statistical difference in that of children (**Figure 4b**).

**Figure 4.**
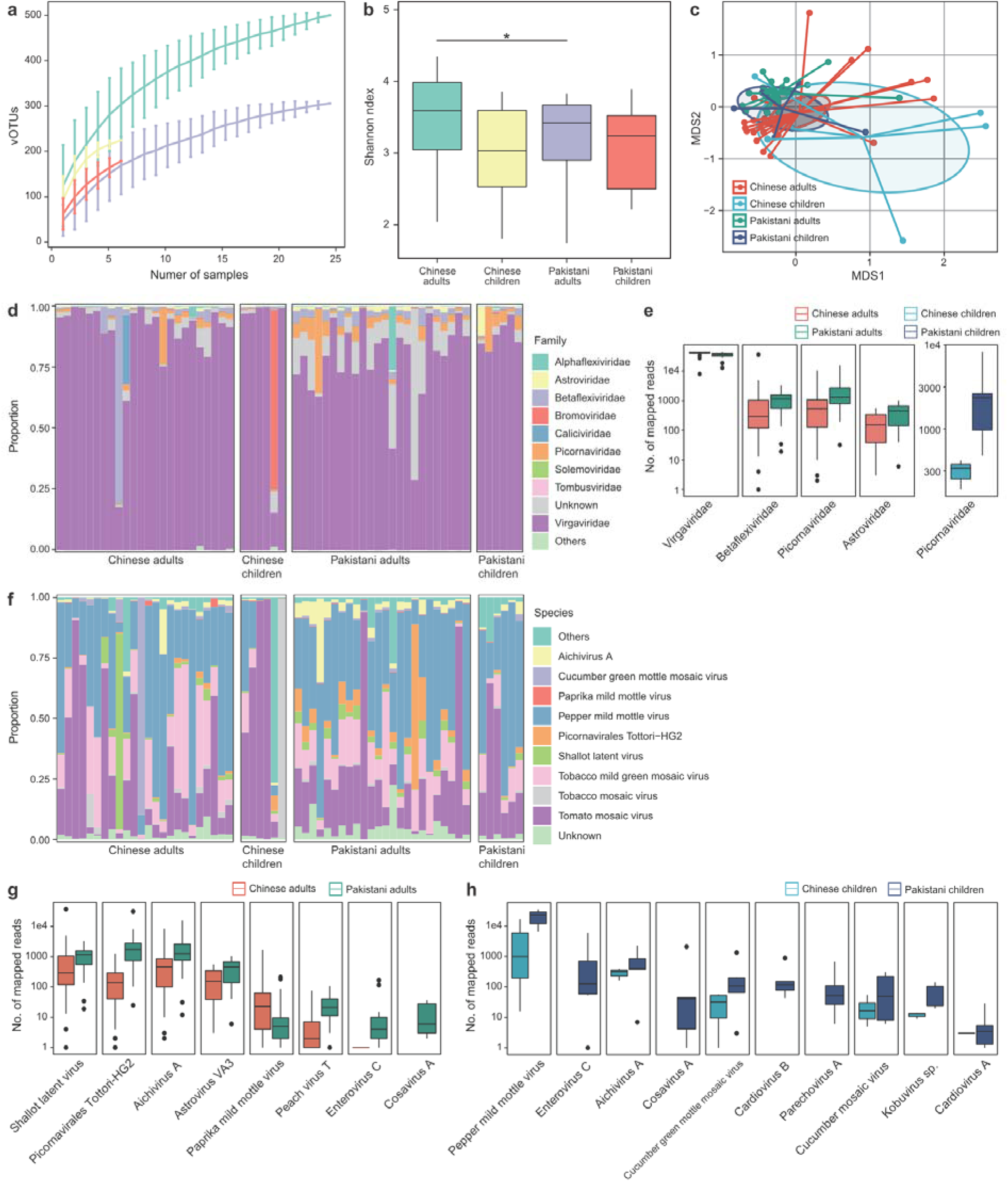
Differences in gut RNA virome between Chinese and Pakistanis. **a**, Rarefaction curve analysis of number of vOTUs on each group of samples. The number of identified vOTUs in different groups is calculated based on a randomly selected specific number of samples with 30 replacements, and the median and quartiles numbers are plotted. **b**, Boxplot shows the Shannon diversity index among four groups. The significance level in the Student’s t test is denoted as: *, *q*<0.05; **, *q*<0.01. c, NMDS analysis based on the composition of virome, revealing the separations between different groups. The location of samples (represented by nodes) in the first two multidimensional scales are shown. Lines connect samples in the same group, and circles cover samples near the center of gravity for each group. **d**, Composition of gut virome at the family level. **e**, Boxplot shows the differential viral families between Chinese and Pakistanis. **f**, Composition of gut virome at the species level. **g-h**, Boxplot shows the differential viral families of adults **(g)** and children **(h)** when compared between Chinese and Pakistanis. For boxplot, boxes represent the interquartile range between the first and third quartiles and median (internal line); whiskers denote the lowest and highest values within 1.5 times the range of the first and third quartiles, respectively; and nodes represent outliers beyond the whiskers.

NMDS analysis on the overall RNA vOTUs composition captured significant separation of adults between Chinese and Pakistanis (*adonis p*<0.001; **Figure 4c**), but of children the separation was visible but not significant (*adonis p*=0.2). Likewise, the viral communities of Chinese adults and children were closer, yet of Pakistani adults and children.

Finally, to investigate the gut RNA viral signatures between Chinese and Pakistanis, we compared two cohorts on viral composition. At the family level, the dominant family *Virgaviridae* consisted of average 83.7% relative abundance in all samples (**Figure 4d**), which was slightly but significantly enriched in Chinese adults compared with that in Pakistani adults (**Figure 4e**). Three other families, *Betaflexiviridae, Picornaviridae*, and *Astroviridae*, was reduced in Chinese adults than in Pakistani adults (Mann-Whitney U test, *q*<0.05 for all), while *Picornaviridae* was also reduced in Chinese children than in Pakistani children. At the species level, the plant-associated virus, including *Pepper mild mottle virus* (average relative abundance, 37.5 ± 23.1%), *Tomato mosaic virus* (27.1 ± 27.4%), and *Tobacco mild green mosaic virus* (14.1 ± 12.4%), composed of the dominant species in all samples (**Figure 4f**). Compared with the Chinese adults, the viral communities of the Pakistani adults showed a significant increase of *Shallot latent virus, Picornavirales Tottori-HG2, Aichivirus A*, and *Astrovirus VA3*, and a remarkable depletion of *Paprika mild mottle virus, Peach virus T, Enterovirus C*, and *Cosavirus A* (**Figure 4g**). When compared with the Chinese children, 9 species were significantly higher in viral communities of Pakistani children (**Figure 4h**), with no species that was lower.

### Concordance between DNA and RNA viromes

Having characterized the differences of DNA and RNA viromes between local Chinese residents and visiting Pakistanis, we wanted to examine the existence of concordance between DNA and RNA viromes. Although the DNA and RNA viromes were irrelevant in Shannon diversity index (Pearson *r*=0.04, *p*=0.7; **Figure 5a**), the overall compositions of two types of viral community were strongly correlated (Procrustes correlation M^2^= 0.37, *p*<0.001; **Figure 5b**). And this correlation was reproducible across nationality and age. Moreover, we identified 24 co-abundance correlations between 6 DNA and 9 RNA viral families (Spearman correlation test *q*<0.05; **Figure 5c**), including some positive correlations between *Adenoviridae* and several RNA viruses and a negative correlation between *Herpesviridae* and *Tombusviridae*. The significance of these relationships required further studies.

**Figure 5.**
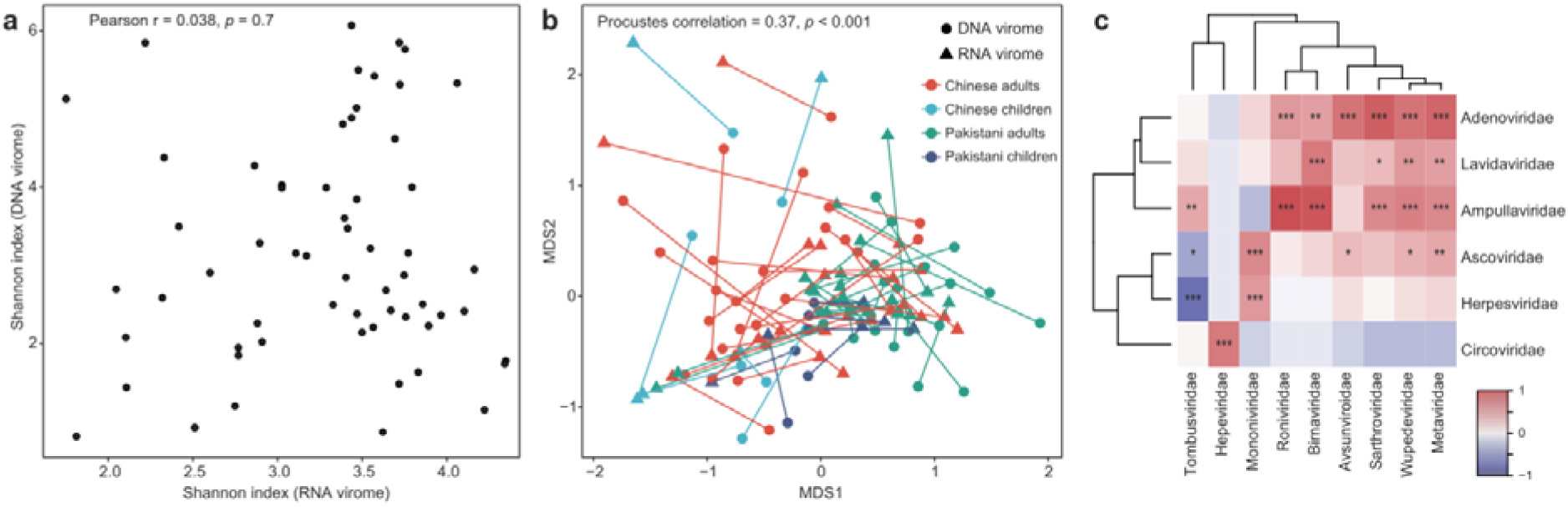
Correlations between DNA and RNA viromes. **a**, Relationship of microdiversity between DNA and RNA virome. **b**, Procrustes analysis of DNA virome versus RNA viromes. Samples for DNA and RNA viromes are shown as circles and blue triangles, respectively; and samples from the same individual are connected by lines. Colors represent samples belong to different groups. **c**, Heatmap shows the co-abundance correlations between DNA and RNA viral families. The significance level in the Spearman correlation test is denoted as: *, *q*<0.05; **, *q*<0.01; ***, *q*<0.001.

### Comparison of bacterial microbiome

For bacterial microbiome, we obtained a total of 1,236 million reads (20.6 ± 7.7 million per sample) from the samples and quantified the relative abundances of a total of 833 taxa, including 12 phyla, 22 classified, 41 orders, 81 families, 179 genera, and 498 species, using MetaPhlAn2 [39]. Comparison on Shannon index showed that the bacterial microbiome of Chinese adults exhibited a significantly higher diversity than that of the Pakistanis (**Figure 6a**), similarly but not significantly trend was observed in that of children. NMDS analysis on the overall bacterial composition also revealed significant separation between Chinese and Pakistan adults (*adonis p*<0.001; **Figure 6b**), as well as between Chinese and Pakistan children (*adonis p*<0.001). Consistent with the observations in DNA and RNA viromes, the bacterial microbiome of Pakistan children was also close to that of Chinese subjects in tendency.

**Figure 6.**
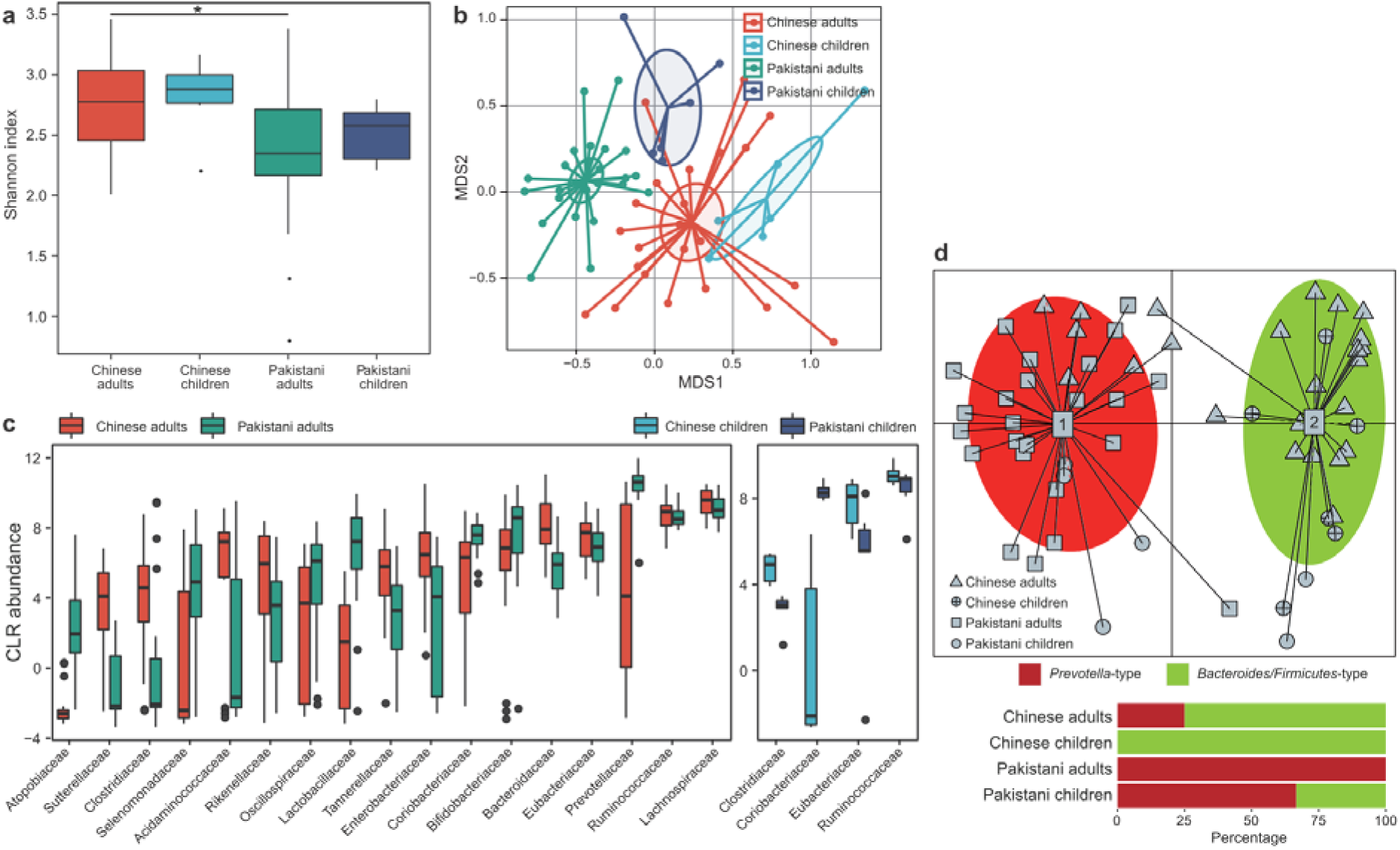
Differences in gut bacterial microbiome between Chinese and Pakistanis. **a**, Boxplot shows the Shannon diversity index among four groups. The significance level in the Student’s t test is denoted as: *, *q*<0.05; **, *q*<0.01. **d**, NMDS analysis based on the composition of bacterial microbiome, revealing the separations between different groups. The location of samples (represented by nodes) in the first two multidimensional scales are shown. Lines connect samples in the same group, and circles cover samples near the center of gravity for each group. **c**, Boxplot shows the bacterial families that differed in abundance between two cohorts. Boxes represent the interquartile range between the first and third quartiles and median (internal line); whiskers denote the lowest and highest values within 1.5 times the range of the first and third quartiles, respectively; and nodes represent outliers beyond the whiskers. **d**, Enterotype analysis of bacterial microbiome samples. The upper panel show the principal component analysis (PCA) of all samples, revealing the separation between two enterotypes. The lower panel show the composition of enterotypes in four groups.

Taxonomically, the bacterial microbiome of Chinese adults showed significant enrichment of *Lachnospiraceae, Ruminococcaceae, Eubacteriaceae, Enterobacteriaceae, Tannerellaceae, Rikenellaceae, Acidaminococcaceae Clostridiaceae*, and *Sutterellaceae* and depletion of *Prevotellaceae, Bifidobacteriaceae, Coriobacteriaceae, Lactobacillaceae, Oscillospiraceae, Selenomonadaceae*, and *Atopobiaceae*, compared with that of Pakistani adults (linear discriminant analysis [LDA] score >3; **Figure 6c**). Similarly, *Clostridiaceae, Eubacteriaceae*, and *Ruminococcaceae* were enriched in Chinese children compared to Pakistani children, and *Coriobacteriaceae* was depleted. At the species level, the Chinese adults exhibited 28 enriched bacterial species and 19 decreased species when compared with the Pakistani adults, while the Chinese children showed 11 enriched species and 12 decreased species compared with the Pakistani children (**Table S3**). The exhibition of enormous differential taxa led to a dramatic distinction of enterotype constitution between Chinese and Pakistanis. The Chinese subjects was characterized by a high proportion of *Bacteroides/Firmicutes*-type (75% and 100% in adults and children, respectively), whereas almost of all Pakistani subjects were *Prevotella*-type (100% in adults and 66.7% in children) (**Figure 6d**).

### Virus-bacteria associations

To study the virus-bacteria correlation, first, we predicted the bacterial hosts of virus by searching the potential viral CRISPR spacers from bacterial metagenomic assemblies (see Methods). This approach allowed host assignments for 3,948 DNA and 4 RNA vOTUs, representing 7.2% and 0.7% of all DNA and RNA viruses, respectively. We revealed a large connection network of family-level known virus (n = 392) and its bacterial host (**Figure 7a**), facilitated by frequent acquisition of phage/prophage in bacterial genomes and spread of phages across bacterial hosts. Members of *Faecalibacterium, Prevotella, Ruminococcus, Bifidobacterium, Dialister*, and *Streptococcus* were the most common host for human gut virome. Meanwhile, the *crAss-like* phages had infected the highest number of bacteria.

**Figure 7.**
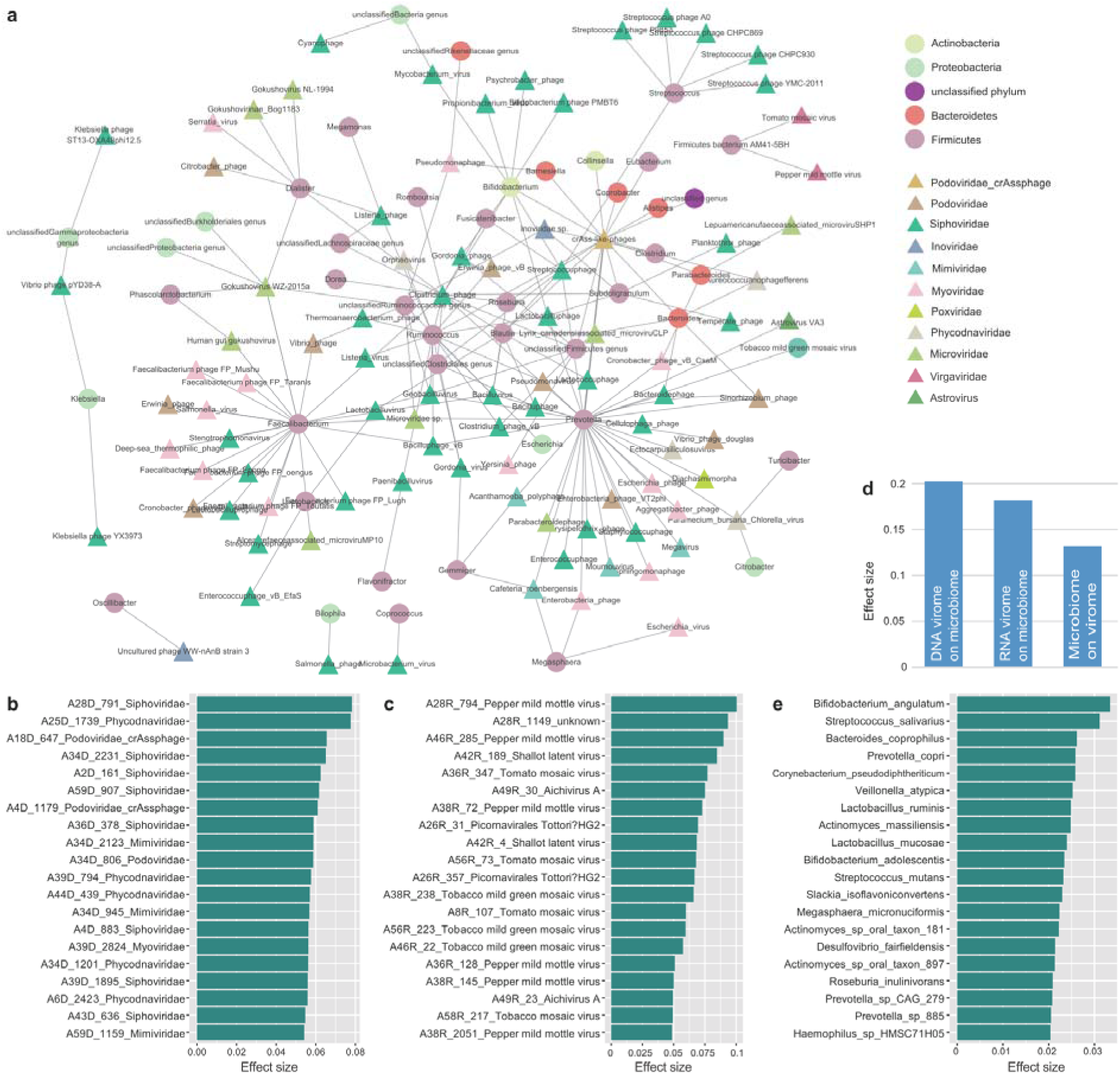
Associations between virome and bacterial microbiome. **a**, Host range of viruses predicted through CRISPR spacer matches. Circles and tringles represent the bacteria and viruses, respectively; and the colors represent their taxonomic assignment at the phylum (for bacteria) or family (for viruses) levels. **b-c**, The 20 DNA **(b)** and RNA families **(c)** for which the highest effect size that significant impact the bacterial microbiome communities. **d**, The combined effect size of viruses on bacterial microbiome as well as bacteria on virome. To calculate the combined effect size, a set of non-redundant covariates (DNA vOTUs, RNA vOTUs, or bacterial species) is selected from the omic datasets, and then the accumulated effect size is calculated by *adonis* analysis using these selected covariates. **e**, The 20 bacterial species with highest effect size for impacting the viral communities.

Then, we performed the PERMANOVA-based effect size analysis between gut virome and microbiome. 287 DNA vOTUs (*q*<0.10), including members of *Siphoviridae, Phycodnaviridae*, and *Podoviridae*-crAssphage, and 25 RNA vOTUs (*q*<0.10) showed significant affection on the bacterial microbiome communities (**Figure 7b-c**). More importantly, combination of these DNA and RNA vOTUs explained 20.2% and 18.2% of the microbiome variance, respectively (**Figure 7d**), suggesting that the effect size of the gut virome on bacterial microbiome is considerable. Similar effect sizes were found in subjects from two nations. Parallelly, 117 bacterial species were identified that significantly impact the holistic composition of DNA and RNA viromes, accounting for 13.2% virome variance (**Figure 7d**). These species included *Bifidobacterium angulatum, Streptococcus salivarius, Bacteroides coprophilus*, and *Prevotella copri* (**Figure 7e**).

## Discussion

Both ethnic origin and residential environment have negligible effects on individual’s gut microbiome [32, 40-42]. To extend this finding on gut virome, our study focused on the viral community of a cohort of Chinese and visiting Pakistanis. Despite sharing the residential environment, the viral diversity and composition of Chinese and Pakistanis were dramatically differed, suggesting that the ethnicity-specific characteristics of virome enable to maintain over an extended period (average 11 and 9 months for Pakistani adults and children, respectively). This result was in accordance with an earlier study showing that the individual characteristics of gut virome can be relatively stable for at least one year [14].

Using *de novo* assembly and discovery approaches, we identified a huge number of viruses from the subjects’ fecal samples, including approximately 55,000 non-redundant complete and partial DNA viral genomes and 569 non-redundant RNA viruses, particularly the number of DNA vOTUs increased over 8-fold compared with the isolated viral sequences in RefSeq database. The majority of viruses were unclassified even at the family level, in agreement with previous observations of extensive novelty of viral world in multiple environments as well as in human gut [43-45].

The DNA viral macrodiversity of Chinese adults was lower than that of Pakistani adults, whereas an opposite phenomenon was observed in the diversity of bacterial community. This result was in conflict with the observation in US adults which exhibited strong correlation between gut virome and microbiome diversities [4]. As most of the DNA viruses were bacteriophages (in this study, the bacterial hosts of at least 7.2% DNA viruses were verified) [4], the degree to which bacterial microbiome drives the virome diversity is considerable. The explanation for high DNA viral diversity in Pakistani adults was unknown, but reason for the enrichment of some eukaryotic viruses in their gut was speculated (see the following discussion). In contrast to DNA virome, the RNA viral diversity was higher in Chinese adults than in Pakistani adults. This observation could be due to the difference of dietary habits between two groups, as in fact the gut RNA viruses were generally plant-associated viruses in our cohort.

Significant compositional differences were observed in DNA and RNA viromes, so was bacterial microbiome between Chinese residents and visiting Pakistanis. In DNA virome, the Pakistani adults showed remarkable enrichment of two eukaryotic viruses, *Adenoviridae* and *Anelloviridae*. Members of *Adenoviridae* were the most prevalent human-associated viruses that can cause respiratory infection, gastroenteritis, and multi-organ diseases [46-48]; while some members of *Anelloviridae* were also associated with human viral infections [49]. *Adenoviridae* was also highly abundant in the gut of Pakistani children but was rare in that of Chinese children, suggesting potential transmission of such viruses from Pakistani parents to their offsprings. In RNA virome, some members of the plant-associated virus *Virgaviridae* were enriched in Pakistanis but some others were reduced. This finding was thought to be connected to the difference of dietary habits between two cohorts. For example, the abundance of *Shallot latent virus* was higher in Pakistani adults than in Chinese adults, as the shallot (e.g. onion, leek) is commonly used in halal foods in the school canteen but rarely appeared in Chinese foods (based on the authors’ experience). In addition, some members of the Pakistani adult-enriched *Picornaviridae*, including *Picornavirales Tottori-HG2, Enterovirus C*, and *Cosavirus A*, and *Astroviridae* were well-known human enteroviruses that can cause diarrhea and enteric infections [50-52]. In bacterial microbiome, the enterotype distribution of Chinese and Pakistanis was deviated, characterized by a high proportion of *Bacteroides/Firmicutes*-type (associated with diets enriched animal carbohydrates [53, 54]) and low proportion of *Prevotella*-type (associated with plant fiber-enriched diets [55]) in Chinese subjects. Combination of these findings suggested that the dietary habits may be a key driver for shaping the gut RNA virome and bacterial microbiome. Of course, more proof-of-principle studies are needed in future.

One striking observation was that the DNA virome of Pakistan children is closer to that of Chinese subjects, when compared with the degree of deviation between Chinese and Pakistan adults. This phenomenon was also observed in RNA virome and bacterial microbiomes in tendency. These findings suggested that the virome and microbiome of children was more changeable than that of adults, despite the fact that the Pakistan adult participants seemed to live a bit longer in China. In accordance with the previous studies, the infant or child gut microbiome was less stable under the changes of environmental, dietary pattern, and antibiotic usage [56-58]. In addition, dynamic development of the infant gut virome towards a more stable adult-like gut virome was also confirmed by recent studies [21, 59, 60].

We characterized the functional capacity of gut virome by identifying over 53,000 KEGG annotated protein-coding genes, of which the core functions seemed consistent with previous findings in the gut phage catalog [61]. Different from the observation in DNA viral composition, the Chinese adults revealed a more diver functional profile than that of the Pakistani adults, as revealed by more metabolism-associated genes in Chinese adults. In addition to general functions, we also identified over 11,000 CAZymes and 37 antibiotic resistance genes from all DNA viruses. To the best of our knowledge, the appearance of extensive CAZymes in gut virome was first found in this study. Potential viral contributions to complex carbon degradation were validated in ocean and soil ecosystems [62, 63]. Thus, our findings further highlight the importance of viral carbohydrate metabolism capacity in human gut. Moreover, the virus-encoded ARGs was also directly relevant to human health, consistent with previous studies [64].

Not only bacteriophages but also free-living viruses in human gut can influence bacterial microbiome structure and therefore indirectly affect health status [65, 66]. We confirmed remarkable connections between viruses and bacterial hosts in our study cohort, including the previous-known parasitic relations (e.g. *crAss-like* phages and Bacteroidetes members [18, 67]) and many novel connections. Noticeably, the Pakistani-dominated genus *Prevotella* connected the largest number of viruses and was responsible for a large part of variance in the virome composition, in agreement with the previous studies showing that the high relative level of *Prevotella* lead to a higher prevalence of temperate bacteriophages and increased virome macrodiversity [14]. One the other hand, we also statistically revealed that the gut virome was also an important determinator of the bacterial microbiome.

As all participants shared the residential environment, we were only able to study the effect of nationality on their gut virome. Through collecting samples from the visiting Pakistani before they arrived China or from other local Pakistani residents, future research is believed to confirm the effect of environment on gut virome. Other limitations in this study included 1) the relatively small sample size, 2) the lack of longitudinal sampling for the individuals, and 3) the inadequacy of viral reference database. These limitations did not affect the robustness of results in the current cohort, but follow-up studies in wider populations will still complement some deficiencies of the current study and provide more new findings.

## Summary

In conclusion, we systematically described the baseline gut virome in a well-characterized cohort of Chinese and visiting Pakistanis and demonstrated that the national background contributed a primary variation to gut virome. The mechanisms underlying the difference between two cohorts remain unclear, but the ethnic factor must be proposed and considered in designing future studies of the virome.

## Methods

### Subject and sample collection

This study received approval from the ethics committee of Dalian Medical University, and written informed consent was obtained from each participant. The methods were carried out in accordance with the approved guidelines. Thirty healthy Pakistani from Dalian Medical University and thirty BMI-, dietary habit-, alcohol intake-and frequency of smoking-matched Chinese healthy controls were recruited for this study. Each cohort was consisted of 24 healthy adults and 6 of their healthy child offsprings. Fresh fecal samples were collected from each subject and were immediately stored at a −80D freezer.

### Experimental procedures for DNA and RNA viromes

#### Virus-like particles enrichment

The procedure of VLPs enrichment was performed on ice. Add 0.1g fecal sample into 1 ml HBSS buffer (137 mM NaCl, 5.4 mM KCl, 1.3 mM CaCl_2_, 0.3 mM Na_2_HPO_4_·2H_2_O, 0.5 mM MgCl_2_·7H_2_O, 0.4 mM KH_2_PO_4_, 0.6 mM MgSO_4_·7H_2_O, 4.2 mM NaHCO_3_, 5.6 mM D-glucose), centrifuge at 10000 g twice to obtain supernatant. After filtering to sterilize, the sterilized filtrate was mixed with the same volume of HBSS buffer and centrifugated at 750,000 g for an hour, the supernatant was stored at −80D. The pellet was collected for DNA extraction.

#### Viral DNA and RNA extraction

The DNA and RNA of virus were extracted by using TIANamp Virus DNA / RNA Kit (TIANGEN) according to the manufacturer’s protocols. Prepare the mixture contained extracted viral DNA, 1μl 20 mM random primers D2-8N (5 ‘-AAGCTAAGACGGCGGTTCGGNNNNNNNN-3’), 1 μl 10xRT mix, 1 μl 10 mM dNTP and 11.5 μl DEPC H_2_O. To synthesize the first strand of viral DNA, desaturated mixture at 95 D for 5 min, add Klenow fragment solution (0.15 μl 10x Klenow Buffer, 0.5 μl Klenow fragment, 0.85 μl DEPC H_2_O) at 37 D. The procedure should be performed twice to obtain two-strand viral DNA. The extracted RNA was reverse transcribed by using Vazyme HiScript II 1st Strand cDNA Synthesis Kit (+gDNA wiper) with the same random amplification primer. The two-strand of cDNA could be synthesized by the same approach.

#### cDNA preparation

Add the mixture contained rSAP and exonuclease-1 into viral two-strand DNA and cDNA at 37 °C, respectively, to remove the remained dNTP and primer D2-8N. After 1 hour, add 10 μl 5X Q5 Reaction Buffer, 3 μl 50 mM MgCl_2_, 1.5 μl 10 mM dNTP, 3 μl 20 mM primer D2 (5 ‘- AAGCTAAGACGGCGGTTCGG -3’), 1.25 μl Q5 High-Fidelity DNA Polymerase and 23.25 μl DEPC H_2_O to amplify the viral DNA and cDNA by polymerase chain reaction (PCR). DNA and cDNA were stored at –20 °C freezer. The DNA and RNA concentration and purity were quantified with NanoDrop2000. DNA and cDNA quality were examined with a 1% agarose gel electrophoresis system.

#### Shotgun sequencing of viromes

All the DNA and cDNA viral samples were subjected to shotgun metagenomic sequencing by using the Illumina HiSeq 3000 platform. Libraries were prepared with a fragment length of approximately 350 bp. Paired-end reads were generated using 150 bp in the forward and reverse directions.

### Bioinformatic analysis of DNA and RNA viromes

#### DNA virome assembly, identification, clustering and taxonomy

The quality control of DNA virome sequences was performed using fastp [68], and the human reads were removed based on Bowtie2 [69] alignment. Each sample was individually assembled using metaSPAdes [70]. Proteins of the contigs were predicted using Prodigal [71]. After that, the assembled contigs (>1,000 bp) were identified as viruses when it satisfied one of the following criteria: 1) at least 3 proteins of a contig (or at least 50% proteins if the contig had less than 6 proteins) were assigned into the viral protein database integrating from NCBI reference viral genomes and the virus orthologous groups database (http://vogdb.org), with a maximum pairwise alignment e-value 1e-10 based on DIAMOND [72]; 2) score >0.7 and *p*-value <0.05 in the VirFinder [73], a k-mer based tool for identifying viral sequences from assembled metagenomic data; 3) at least 2 proteins were uncharacterized from the integrated databases of KEGG [36], NCBI-nr, and UniProt [74]. Viral contigs were pairwise blasted and the highly consistent viruses with 95% nucleotide identity and 80% coverage of the sequence were further clustered into vOTUs using inhouse scripts. The longest viral contig was defined as representative sequence for each vOTU. Proteins of the vOTUs were aligned with the available viral proteins using blastp (minimum score 50), and the family level taxonomy of a vOTU was generated if more than a third of its proteins were assigned into the same viral family.

#### Macrodiversity and microdiversity of DNA virome

The macrodiversity (Shannon diversity index) of virome was calculated using *vegan* package in R platform, with a uniformed number of reads (1 million) for each sample. The microdiversity (nucleotide diversity, π) for representative sequence in each vOTU was calculated based on the methodology developed by Schloissnig *et al*. [75], and microdiversity of a sample was generated by averaging from the viruses that presented (depth >10x) in that sample.

#### Functional profiles of DNA virome

The viral proteins were aligned to KEGG [36] database (blastp similarity >30%) for functional annotation. For functional profiling, the KEGG aligned proteins were dereplicated with CD-HIT [76] (>95% identity and >90% sequence coverage) to construct the custom viral functional gene catalog, followed by mapping the reads to the catalog using the ‘very-sensitive-local’ setting in Bowtie2 [69]. The relative abundance of each functional gene in sample was normalized by the total numbers of viral reads (the reads mapped to the viral sequence) in the sample, and was transformed into centered log ratio (CLR) coordinates using *microbiome* package in R platform. The carbohydrate-active enzymes and acquired antibiotic resistance genes for the viruses were predicted from the CAZy [37] and CARD [77] databases, respectively, using the same manner as functional assignment.

#### RNA viromes assembly, identification, clustering and taxonomy

The metatranscriptomic data of RNA virome reads was trimmed using fastp [68]. The contamination of ribosomal RNA reads was identified and removed by mapping to the small subunit sequences (bacterial 16S and eukaryotic 18S) on the latest SILVA database [78]. The rnaSPAdes was utilized in metatranscriptomic assembly for each sample [79]. To identify RNA viruses, the assembled contigs (>500 bp) was aligned to the reference RNA virus proteins downloaded from GenBank database using DIAMOND (blastx e-value <1e-5). We also identified the RNA viral contigs by searching the RNA-dependent RNA polymerase genes (RdRp genes, referred from Evan *et al*. [80]) using a Hidden Markov Model approach [81]. Then, the RNA viral sequences were clustered based on 95% identity and 90% coverage of the sequence.

### Bacterial microbiome sequencing and analysis

All raw metagenomic data was trimmed and the human contamination sequences was removed using the same methods in virome. MetaPhlan2 [39] was employed to generate the taxonomic profile for each sample using default parameters. Enterotype analysis was performed at the bacterial genus level composition based on the methodology developed by Costea *et al*. [55]. The high quality microbiome data was assembled using metaSPAdes [70], and the resulting contigs was searched against the NCBI-nt database to identity the bacteria sequence (>70% similarity and >70% coverage at the phylum level). To search the potential bacterial host of virus, the CRISPR spacers in bacteria sequence was predicted using PILER-CR [82], and then the spacers were blasted to the viral sequences (‘‘blastn-short’’ mode and bitscore >50) to identify the phage-bacterial host pairs. The matching bacterial host and viral sequence was summarized at the genus level. To avoid ambiguity, genus producing highest number of spacers hits was considered as primary host.

### Statistical analysis

Statistical analyses were implemented at the R 3.6 platform (https://www.r-project.org/). Permutational multivariate analysis of variance (PERMANOVA) was performed with the *adonis* function of the *vegan* package, and the *adonis P*-value was generated based on 1,000 permutations. The method of effect size analysis was referred as Wang *et al*. [10]. The no-metric multidimensional scaling (NMDS) analysis was used as the ordination methods (*metaMDS* function in *vegan* package) for compositional data. The Procrustes coordinates analysis and significance were generated using the *procuste* and *procuste*.*randtest* functions in *vegan* package. The principal component analysis (PCA) was performed and visualized using the *ade4* package. The Wilcoxon rank-sum test was used to measure statistical differences in diversity and taxonomic levels between two cohorts. *P*-values were corrected for multiple testing using the Benjamini-Hochberg procedure.

## Data availability

The raw sequencing dataset acquired in this study has been deposited to the NCBI SRA database under the accession code PRJNA641593. The sample metadata, vOTU and taxonomic composition data, and the statistical scripts are available from the corresponding author on reasonable request.

## Acknowledgements

This work was supported by grants from the Priority Academic Program Development of Jiangsu Higher Education Institutions (Integration of Chinese and Western Medicine), the National Natural Science Foundation of China (No. 81902037) and the Liaoning Provincial Natural Science Foundation (No. 20180530086).

## Author contributions

T. M., S. L., Y. M., and Q.Y. conceived and directed the study. Q. Y., Y. W., X. C., G. W., T. A. and X. L. developed and conducted the experiments. Q. Y., G. W. and T. A. performed sample collection and investigation. H. J., K. G., Y. Z., and P. Z. carried out data processing and analyses. S. L., Q. Y., and T. M. drafted the manuscript. Y. M., G. W., Y. L.; J. W.; G. C.; A. Z. and P. L. participated in design and coordination, and helped draft the manuscript. P. Z., Y. S., M. X. and P. L. revised the manuscript. All authors read and approved the final manuscript.

---

**Supplementary figure 1.**
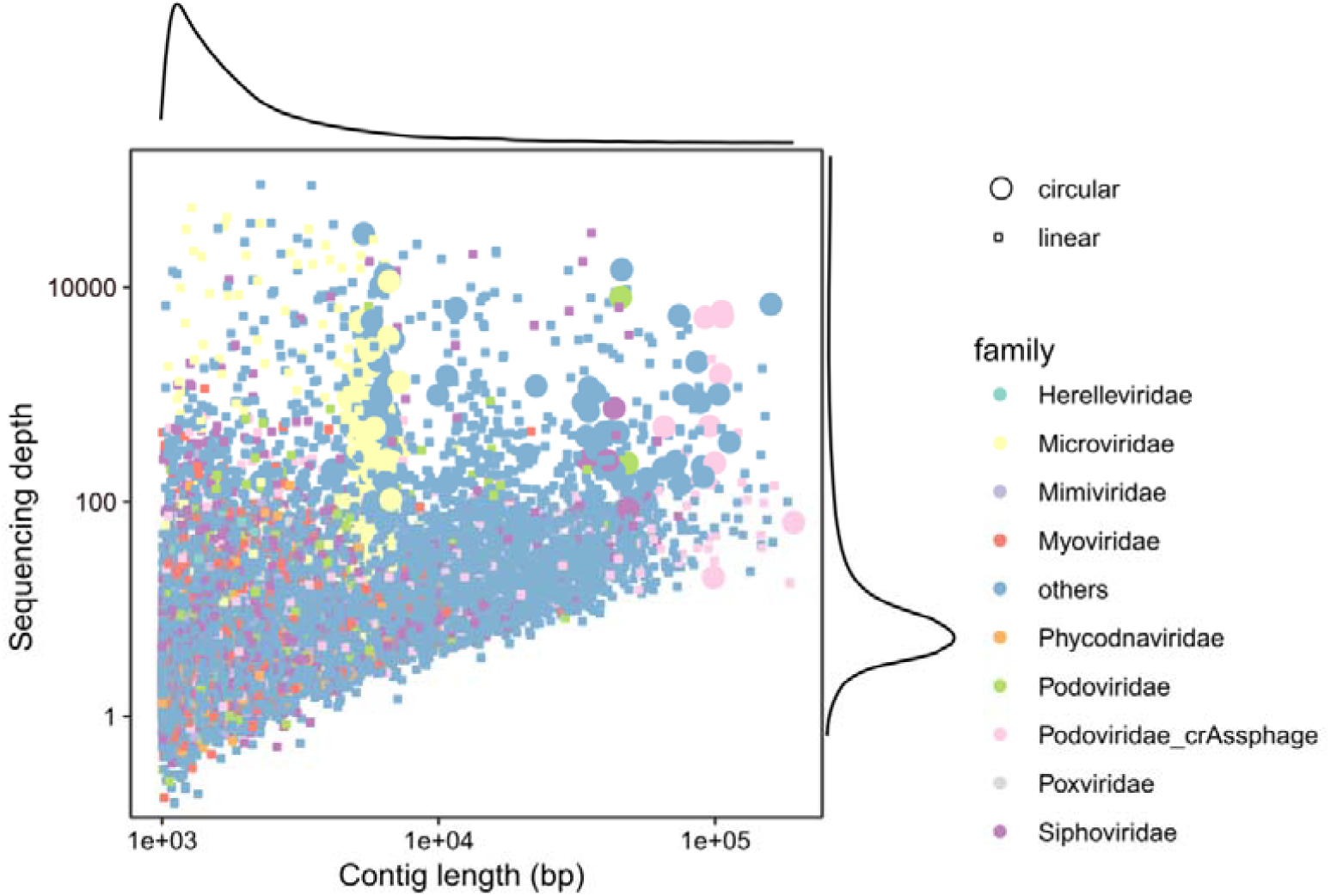
Distribution of DNA viral contigs by length and depth of coverage.

## Reference

1. Sender R, Fuchs S, Milo R. Are We Really Vastly Outnumbered? Revisiting the Ratio of Bacterial to Host Cells in Humans. Cell. 2016; 164(3):337–340.

2. Vandeputte D, Kathagen G, D’Hoe K, Vieira-Silva S, Valles-Colomer M, Sabino J, Wang J, Tito RY, De Commer L, Darzi Y et al. Quantitative microbiome profiling links gut community variation to microbial load. Nature. 2017; 551(7681):507–511.

3. Castro-Mejia JL, Muhammed MK, Kot W, Neve H, Franz CM, Hansen LH, Vogensen FK, Nielsen DS. Optimizing protocols for extraction of bacteriophages prior to metagenomic analyses of phage communities in the human gut. Microbiome. 2015; 3:64.

4. Moreno-Gallego JL, Chou SP, Di Rienzi SC, Goodrich JK, Spector TD, Bell JT, Youngblut ND, Hewson I, Reyes A, Ley RE. Virome Diversity Correlates with Intestinal Microbiome Diversity in Adult Monozygotic Twins. Cell Host Microbe. 2019; 25(2):261–272 e265.

5. Marchesi JR. Prokaryotic and eukaryotic diversity of the human gut. Adv Appl Microbiol. 2010; 72:43–62.

6. Turnbaugh PJ, Ley RE, Mahowald MA, Magrini V, Mardis ER, Gordon JI. An obesity-associated gut microbiome with increased capacity for energy harvest. Nature. 2006; 444(7122):1027–1031.

7. Qin J, Li R, Raes J, Arumugam M, Burgdorf KS, Manichanh C, Nielsen T, Pons N, Levenez F, Yamada T et al. A human gut microbial gene catalogue established by metagenomic sequencing. Nature. 2010; 464(7285):59–65.

8. Lloyd-Price J, Mahurkar A, Rahnavard G, Crabtree J, Orvis J, Hall AB, Brady A, Creasy HH, McCracken C, Giglio MG et al. Strains, functions and dynamics in the expanded Human Microbiome Project. Nature. 2017; 550(7674):61–66.

9. Pedersen HK, Gudmundsdottir V, Nielsen HB, Hyotylainen T, Nielsen T, Jensen BA, Forslund K, Hildebrand F, Prifti E, Falony G et al. Human gut microbes impact host serum metabolome and insulin sensitivity. Nature. 2016; 535(7612):376–381.

10. Wang X, Yang S, Li S, Zhao L, Hao Y, Qin J, Zhang L, Zhang C, Bian W, Zuo L et al. An aberrant gut microbiota alters host metabolome and impacts renal failure in human and rodents. 2020. doi:10.1136/gutjnl-2019-319766.

11. Qin J, Li Y, Cai Z, Li S, Zhu J, Zhang F, Liang S, Zhang W, Guan Y, Shen D et al. A metagenome-wide association study of gut microbiota in type 2 diabetes. Nature. 2012; 490(7418):55–60.

12. Quigley EM. Gut bacteria in health and disease. Gastroenterol Hepatol (N Y). 2013; 9(9):560–569.

13. Handley SA. The virome: a missing component of biological interaction networks in health and disease. Genome Med. 2016; 8(1):32.

14. Shkoporov AN, Clooney AG, Sutton TDS, Ryan FJ, Daly KM, Nolan JA, McDonnell SA, Khokhlova EV, Draper LA, Forde A et al. The Human Gut Virome Is Highly Diverse, Stable, and Individual Specific. Cell Host Microbe. 2019; 26(4):527–541 e525.

15. Forster SC, Kumar N, Anonye BO, Almeida A, Viciani E, Stares MD, Dunn M, Mkandawire TT, Zhu A, Shao Y et al. A human gut bacterial genome and culture collection for improved metagenomic analyses. Nat Biotechnol. 2019; 37(2):186–192.

16. Zou Y, Xue W, Luo G, Deng Z, Qin P, Guo R, Sun H, Xia Y, Liang S, Dai Y et al. 1,520 reference genomes from cultivated human gut bacteria enable functional microbiome analyses. Nat Biotechnol. 2019; 37(2):179–185.

17. Norman JM, Handley SA, Baldridge MT, Droit L, Liu CY, Keller BC, Kambal A, Monaco CL, Zhao G, Fleshner P et al. Disease-specific alterations in the enteric virome in inflammatory bowel disease. Cell. 2015; 160(3):447–460.

18. Shkoporov AN, Khokhlova EV, Fitzgerald CB, Stockdale SR, Draper LA, Ross RP, Hill C. PhiCrAss001 represents the most abundant bacteriophage family in the human gut and infects Bacteroides intestinalis. Nat Commun. 2018; 9(1):4781.

19. Yutin N, Makarova KS, Gussow AB, Krupovic M, Segall A, Edwards RA, Koonin EV. Discovery of an expansive bacteriophage family that includes the most abundant viruses from the human gut. Nat Microbiol. 2018; 3(1):38–46.

20. Minot S, Bryson A, Chehoud C, Wu GD, Lewis JD, Bushman FD. Rapid evolution of the human gut virome. Proc Natl Acad Sci U S A. 2013; 110(30):12450–12455.

21. Lim ES, Zhou Y, Zhao G, Bauer IK, Droit L, Ndao IM, Warner BB, Tarr PI, Wang D, Holtz LR. Early life dynamics of the human gut virome and bacterial microbiome in infants. Nat Med. 2015; 21(10):1228–1234.

22. Hoyles L, McCartney AL, Neve H, Gibson GR, Sanderson JD, Heller KJ, van Sinderen D. Characterization of virus-like particles associated with the human faecal and caecal microbiota. Res Microbiol. 2014; 165(10):803–812.

23. Kleiner M, Hooper LV, Duerkop BA. Evaluation of methods to purify virus-like particles for metagenomic sequencing of intestinal viromes. BMC Genomics. 2015; 16:7.

24. Pannaraj PS, Ly M, Cerini C, Saavedra M, Aldrovandi GM, Saboory AA, Johnson KM, Pride DT. Shared and Distinct Features of Human Milk and Infant Stool Viromes. Front Microbiol. 2018; 9:1162.

25. Maqsood R, Rodgers R, Rodriguez C, Handley SA, Ndao IM, Tarr PI, Warner BB, Lim ES, Holtz LR. Discordant transmission of bacteria and viruses from mothers to babies at birth. Microbiome. 2019; 7(1):156.

26. Nakatsu G, Zhou H, Wu WKK, Wong SH, Coker OO, Dai Z, Li X, Szeto CH, Sugimura N, Lam TY et al. Alterations in Enteric Virome Are Associated With Colorectal Cancer and Survival Outcomes. Gastroenterology. 2018; 155(2):529–541 e525.

27. Hannigan GD, Duhaime MB, Ruffin MTt, Koumpouras CC, Schloss PD. Diagnostic Potential and Interactive Dynamics of the Colorectal Cancer Virome. mBio. 2018; 9(6).

28. Zuo T, Lu XJ, Zhang Y, Cheung CP, Lam S, Zhang F, Tang W, Ching JYL, Zhao R, Chan PKS et al. Gut mucosal virome alterations in ulcerative colitis. Gut. 2019; 68(7):1169–1179.

29. Zhao G, Vatanen T, Droit L, Park A, Kostic AD, Poon TW, Vlamakis H, Siljander H, Harkonen T, Hamalainen AM et al. Intestinal virome changes precede autoimmunity in type I diabetes-susceptible children. Proc Natl Acad Sci U S A. 2017; 114(30):E6166–E6175.

30. Guo L, Hua X, Zhang W, Yang S, Shen Q, Hu H, Li J, Liu Z, Wang X, Wang H et al. Viral metagenomics analysis of feces from coronary heart disease patients reveals the genetic diversity of the Microviridae. Virol Sin. 2017; 32(2):130–138.

31. Vangay P, Johnson AJ, Ward TL, Al-Ghalith GA, Shields-Cutler RR, Hillmann BM, Lucas SK, Beura LK, Thompson EA, Till LM et al. US Immigration Westernizes the Human Gut Microbiome. Cell. 2018; 175(4):962–972 e910.

32. Deschasaux M, Bouter KE, Prodan A, Levin E, Groen AK, Herrema H, Tremaroli V, Bakker GJ, Attaye I, Pinto-Sietsma SJ et al. Depicting the composition of gut microbiota in a population with varied ethnic origins but shared geography. Nat Med. 2018; 24(10):1526–1531.

33. He Y, Wu W, Zheng HM, Li P, McDonald D, Sheng HF, Chen MX, Chen ZH, Ji GY, Zheng ZD et al. Regional variation limits applications of healthy gut microbiome reference ranges and disease models. Nat Med. 2018; 24(10):1532–1535.

34. Sun J, Liao XP, D’Souza AW, Boolchandani M, Li SH, Cheng K, Luis Martinez J, Li L, Feng YJ, Fang LX et al. Environmental remodeling of human gut microbiota and antibiotic resistome in livestock farms. Nat Commun. 2020; 11(1):1427.

35. Gregory AC, Zayed AA, Conceicao-Neto N, Temperton B, Bolduc B, Alberti A, Ardyna M, Arkhipova K, Carmichael M, Cruaud C et al. Marine DNA Viral Macro- and Microdiversity from Pole to Pole. Cell. 2019; 177(5):1109–1123 e1114.

36. Kanehisa M, Furumichi M, Tanabe M, Sato Y, Morishima K. KEGG: new perspectives on genomes, pathways, diseases and drugs. Nucleic Acids Res. 2017; 45(D1):D353–D361.

37. Lombard V, Golaconda Ramulu H, Drula E, Coutinho PM, Henrissat B. The carbohydrate-active enzymes database (CAZy) in 2013. Nucleic Acids Res. 2014; 42(Database issue):D490–495.

38. Wolf YI, Kazlauskas D, Iranzo J, Lucia-Sanz A, Kuhn JH, Krupovic M, Dolja VV, Koonin EV. Origins and Evolution of the Global RNA Virome. mBio. 2018;9(6):e02329–18.

39. Truong DT, Franzosa EA, Tickle TL, Scholz M, Weingart G, Pasolli E, Tett A, Huttenhower C, Segata N. MetaPhlAn2 for enhanced metagenomic taxonomic profiling. Nat Methods. 2015; 12(10):902–903.

40. Gupta VK, Paul S, Dutta C. Geography, Ethnicity or Subsistence-Specific Variations in Human Microbiome Composition and Diversity. Front Microbiol. 2017; 8:1162.

41. Gaulke CA, Sharpton TJ. The influence of ethnicity and geography on human gut microbiome composition. Nat Med. 2018; 24(10):1495–1496.

42. Korpela K, Costea P, Coelho LP, Kandels-Lewis S, Willemsen G, Boomsma DI, Segata N, Bork P. Selective maternal seeding and environment shape the human gut microbiome. Genome Res. 2018; 28(4):561–568.

43. Reyes A, Semenkovich NP, Whiteson K, Rohwer F, Gordon JI. Going viral: next-generation sequencing applied to phage populations in the human gut. Nat Rev Microbiol. 2012; 10(9):607–617.

44. Paez-Espino D, Eloe-Fadrosh EA, Pavlopoulos GA, Thomas AD, Huntemann M, Mikhailova N, Rubin E, Ivanova NN, Kyrpides NC. Uncovering Earth’s virome. Nature. 2016; 536(7617):425–430.

45. Rampelli S, Turroni S, Schnorr SL, Soverini M, Quercia S, Barone M, Castagnetti A, Biagi E, Gallinella G, Brigidi P et al. Characterization of the human DNA gut virome across populations with different subsistence strategies and geographical origin. Environ Microbiol. 2017; 19(11):4728–4735.

46. Wadell G. Adenoviridae. the adenoviruses. In: Laboratory Diagnosis of Infectious Diseases Principles and Practice. Springer. 1988: 284–300.

47. Centers for Disease C, Prevention. Acute respiratory disease associated with adenovirus serotype 14--four states, 2006-2007. MMWR Morb Mortal Wkly Rep. 2007; 56(45):1181–1184.

48. Jones MS, Harrach B, Ganac RD, Gozum MM, dela Cruz WP, Riedel B, Pan C, Delwart EL, Schnurr DP. New adenovirus species found in a patient presenting with gastroenteritis. Journal of virology. 2007; 81(11):5978–5984.

49. Bernardin F, Operskalski E, Busch M, Delwart E. Transfusion transmission of highly prevalent commensal human viruses. Transfusion. 2010; 50(11):2474–2483.

50. Johnston S, Sanderson G, Pattemore P, Smith S, Bardin P, Bruce C, Lambden P, Tyrrell D, Holgate S. Use of polymerase chain reaction for diagnosis of picornavirus infection in subjects with and without respiratory symptoms. Journal of clinical microbiology. 1993; 31(1):111–117.

51. Monroe SS, Jiang B, Stine SE, Koopmans M, Glass R. Subgenomic RNA sequence of human astrovirus supports classification of Astroviridae as a new family of RNA viruses. Journal of virology. 1993; 67(6):3611–3614.

52. Rotbart HA. Treatment of picornavirus infections. Antiviral research. 2002; 53(2):83–98.

53. De Filippo C, Cavalieri D, Di Paola M, Ramazzotti M, Poullet JB, Massart S, Collini S, Pieraccini G, Lionetti P. Impact of diet in shaping gut microbiota revealed by a comparative study in children from Europe and rural Africa. Proc Natl Acad Sci U S A. 2010; 107(33):14691–14696.

54. David LA, Maurice CF, Carmody RN, Gootenberg DB, Button JE, Wolfe BE, Ling AV, Devlin AS, Varma Y, Fischbach MA et al. Diet rapidly and reproducibly alters the human gut microbiome. Nature. 2014; 505(7484):559–563.

55. Costea PI, Hildebrand F, Arumugam M, Backhed F, Blaser MJ, Bushman FD, de Vos WM, Ehrlich SD, Fraser CM, Hattori M et al. Enterotypes in the landscape of gut microbial community composition. Nat Microbiol. 2018; 3(1):8–16.

56. Koenig JE, Spor A, Scalfone N, Fricker AD, Stombaugh J, Knight R, Angenent LT, Ley RE. Succession of microbial consortia in the developing infant gut microbiome. Proc Natl Acad Sci U S A. 2011;108 Suppl 1:4578–4585.

57. Yassour M, Vatanen T, Siljander H, Hamalainen AM, Harkonen T, Ryhanen SJ, Franzosa EA, Vlamakis H, Huttenhower C, Gevers D et al. Natural history of the infant gut microbiome and impact of antibiotic treatment on bacterial strain diversity and stability. Sci Transl Med. 2016; 8(343):343ra381.

58. Chu DM, Antony KM, Ma J, Prince AL, Showalter L, Moller M, Aagaard KM. The early infant gut microbiome varies in association with a maternal high-fat diet. Genome Med. 2016;8(1):77.

59. Reyes A, Blanton LV, Cao S, Zhao G, Manary M, Trehan I, Smith MI, Wang D, Virgin HW, Rohwer F et al. Gut DNA viromes of Malawian twins discordant for severe acute malnutrition. Proc Natl Acad Sci U S A. 2015; 112(38):11941–11946.

60. Beller L, Matthijnssens J. What is (not) known about the dynamics of the human gut virome in health and disease. Current opinion in virology. 2019; 37:52–57.

61. Ma Y, You X, Mai G, Tokuyasu T, Liu C. A human gut phage catalog correlates the gut phageome with type 2 diabetes. Microbiome. 2018; 6(1):24.

62. Suttle CA. Marine viruses--major players in the global ecosystem. Nat Rev Microbiol. 2007; 5(10):801–812.

63. Emerson JB, Roux S, Brum JR, Bolduc B, Woodcroft BJ, Jang HB, Singleton CM, Solden LM, Naas AE, Boyd JA et al. Host-linked soil viral ecology along a permafrost thaw gradient. Nat Microbiol. 2018; 3(8):870–880.

64. Ogilvie LA, Bowler LD, Caplin J, Dedi C, Diston D, Cheek E, Taylor H, Ebdon JE, Jones BV. Genome signature-based dissection of human gut metagenomes to extract subliminal viral sequences. Nat Commun. 2013; 4:2420.

65. Scarpellini E, Ianiro G, Attili F, Bassanelli C, De Santis A, Gasbarrini A. The human gut microbiota and virome: Potential therapeutic implications. Dig Liver Dis. 2015; 47(12):1007–1012.

66. Foca A, Liberto MC, Quirino A, Marascio N, Zicca E, Pavia G. Gut inflammation and immunity: what is the role of the human gut virome? Mediators Inflamm. 2015; 2015:326032.

67. Guerin E, Shkoporov A, Stockdale SR, Clooney AG, Ryan FJ, Sutton TDS, Draper LA, Gonzalez-Tortuero E, Ross RP, Hill C. Biology and Taxonomy of crAss-like Bacteriophages, the Most Abundant Virus in the Human Gut. Cell Host Microbe. 2018; 24(5):653–664 e656.

68. Chen S, Zhou Y, Chen Y, Gu J. fastp: an ultra-fast all-in-one FASTQ preprocessor. Bioinformatics. 2018; 34(17):i884–i890.

69. Langmead B, Salzberg SL. Fast gapped-read alignment with Bowtie 2. Nat Methods. 2012; 9(4):357–359.

70. Nurk S, Meleshko D, Korobeynikov A, Pevzner PA. metaSPAdes: a new versatile metagenomic assembler. Genome Res. 2017; 27(5):824–834.

71. Hyatt D, Chen GL, Locascio PF, Land ML, Larimer FW, Hauser LJ. Prodigal: prokaryotic gene recognition and translation initiation site identification. BMC Bioinformatics. 2010; 11:119.

72. Buchfink B, Xie C, Huson DH. Fast and sensitive protein alignment using DIAMOND. Nat Methods. 2015; 12(1):59–60.

73. Ren J, Ahlgren NA, Lu YY, Fuhrman JA, Sun FZ. VirFinder: a novel k-mer based tool for identifying viral sequences from assembled metagenomic data. Microbiome. 2017; 5.

74. Apweiler R. Activities at the Universal Protein Resource (UniProt) (vol 42, pg D198, 2014). Nucleic Acids Res. 2014; 42(11):7486–7486.

75. Schloissnig S, Arumugam M, Sunagawa S, Mitreva M, Tap J, Zhu A, Waller A, Mende DR, Kultima JR, Martin J et al. Genomic variation landscape of the human gut microbiome. Nature. 2013; 493(7430):45–50.

76. Li W, Godzik A. Cd-hit: a fast program for clustering and comparing large sets of protein or nucleotide sequences. Bioinformatics. 2006; 22(13):1658–1659.

77. Jia B, Raphenya AR, Alcock B, Waglechner N, Guo P, Tsang KK, Lago BA, Dave BM, Pereira S, Sharma AN et al. CARD 2017: expansion and model-centric curation of the comprehensive antibiotic resistance database. Nucleic Acids Res. 2017; 45(D1):D566–D573.

78. Quast C, Pruesse E, Yilmaz P, Gerken J, Schweer T, Yarza P, Peplies J, Glockner FO. The SILVA ribosomal RNA gene database project: improved data processing and web-based tools. Nucleic Acids Res. 2013; 41(Database issue):D590–596.

79. Bushmanova E, Antipov D, Lapidus A, Prjibelski AD. rnaSPAdes: a de novo transcriptome assembler and its application to RNA-Seq data. Gigascience. 2019;8(9):giz100.

80. Starr EP, Nuccio EE, Pett-Ridge J, Banfield JF, Firestone MK. Metatranscriptomic reconstruction reveals RNA viruses with the potential to shape carbon cycling in soil. P Natl Acad Sci USA. 2019; 116(51):25900–25908.

81. Johnson LS, Eddy SR, Portugaly E. Hidden Markov model speed heuristic and iterative HMM search procedure. Bmc Bioinformatics. 2010; 11.

82. Edgar RC. PILER-CR: Fast and accurate identification of CRISPR repeats. Bmc Bioinformatics. 2007; 8.

